# Spatial structure formation by RsmE-regulated extracellular secretions in *Pseudomonas fluorescens* Pf0-1

**DOI:** 10.1101/2022.07.26.501654

**Authors:** Anton Evans, Meghan Wells, Jordan Denk, William Mazza, Raziel Santos, Amber Delprince, Wook Kim

## Abstract

Cells in microbial communities on surfaces live and divide in close proximity, which greatly enhances the potential for social interactions. Spatiogenetic structures manifest through competitive and cooperative interactions among the same and different genotypes within a shared space, and extracellular secretions appear to function dynamically at the forefront. A previous experimental evolution study utilizing *Pseudomonas fluorescens* Pf0-1 colonies demonstrated that diverse mutations in the *rsmE* gene are repeatedly and exclusively selected through the formation of a dominant spatial structure. RsmE’s primary molecular function is translation repression, and its homologs regulate various social and virulence phenotypes. *Pseudomonas* spp. possess multiple paralogs of Rsm proteins, and RsmA, RsmE, and RsmI are the most prevalent. Here, we demonstrate that the production of a mucoid polymer and a biosurfactant are exclusively regulated through RsmE, contradicting the generalized notion of functional redundancy among the Rsm paralogs. Furthermore, we identify the biosurfactant as the cyclic lipopeptide gacamide A. Competition and microscopy analyses show that the mucoid polymer is solely responsible for creating a space of low cellular density, which is shared exclusively by the same genotype. Gacamide A and other RsmE-regulated products appear to establish a physical boundary that prevents the encroachment of the competing genotype into the newly created space. Although cyclic lipopeptides and other biosurfactants are best known for their antimicrobial properties and reducing surface tension to promote the spreading of cells on various surfaces, they also appear to help define spatial structure formation within a dense community.

**IMPORTANCE:** In densely populated colonies of the bacterium *Pseudomonas fluorescens* Pf0-1, diverse mutations in the *rsmE* gene are naturally selected by solving the problem of overcrowding. Here, we show that RsmE-regulated secretions function together to create and protect space of low cell density. A biosurfactant generally promotes the spreading of bacterial cells on abiotic surfaces, however, it appears to function atypically within a crowded population by physically defining genotypic boundaries. Another significant finding is that these secretions are not regulated by RsmE’s paralogs that share high sequence similarity. The experimental pipeline described in this study is highly tractable and should facilitate future studies to explore additional RsmE-regulated products and address why RsmE is functionally unique from its paralogs.

## INTRODUCTION

Central to the architecture of microbial communities is the extracellular matrix [1-8], a dynamic cumulus of compounds produced by individual cells that physically define both the spatial arrangements within and the three-dimensional boundaries. Micron-scale spatiogenetic structures readily emerge within surface-grown communities as individual cells produce identical copies of themselves in a given area [9-11]. Competition between different genotypes lead to the spatial enrichment of a particular genotype, producing macroscopic regions that stem from a recent common ancestor [10, 12-15]. Individual phenotypes could positively or negatively impact the fitness of neighboring cells, including the consumption of limiting nutrients and the secretion of enzymes and toxins that promote or discourage the growth of neighboring cells [8, 16-18]. Mechanistic understanding of how individual phenotypes antagonize or synergize with another clearly carries both fundamental and clinical significance.

Researchers employ various experimental approaches to study the interactive dynamics of microbial cells within a community, whether it be computationally [9, 10, 19] or empirically on a variety of abiotic surfaces [5, 20-22]. We have previously described a model system based on *Pseudomonas fluorescens* colonies, which shows how spatial structures rapidly evolve within clonal aggregates [23]. Mucoid patches repeatedly emerge on the surface of aging colonies due to the activity of specific mutants where they expand space and decrease local density.

Remarkably, a mutation in a single gene, *rsmE*, was responsible for each and every case of over 500 independently derived mucoid patches. Importantly, *rsmE* mutants share the same growth rate in isolation compared to the parent cells, and the evolutionary advantage specifically requires the proximal presence of the parent cells. These observations collectively suggest that RsmE-regulated phenotypes physically act to create dominant spatial structures in a densely populated bacterial colony.

RsmE belongs to the CsrA/Rsm family and its homologs are a regulator of social and virulence phenotypes in gamma-proteobacteria [24, 25]. CsrA was the first member of the family to be discovered three decades ago in *Escherichia coli* [26], and its homologs are now known to be present in over 2900 species [27]. CsrA/Rsm proteins interact with diverse mRNA [25, 28-30] and primarily function as a translation repressor by either directly or indirectly blocking their respective Shine-Dalgarno (SD) sequence [31-34]. CsrA also possesses additional regulatory functions that impact Rho-dependent transcription attenuation, mRNA stabilization and destabilization, and even activation of translation [32]. In contrast to CsrA in Enterobacteriaceae, *Pseudomonas* spp. possess varying numbers of Rsm paralogs [27]. Rsm paralogs were initially characterized to repress the production of secondary metabolites and are generally described to overlap or cumulate in function [35-41].

Although the three paralogs in *P. fluorescens* (RsmE, RsmA, and RsmI) share high sequence similarity, the exclusive selection of mutations in the *rsmE* locus [23] suggests functional specificity of RsmE from its paralogs. Here, we show that all three Rsm paralogs are expressed, but RsmE uniquely governs the production of both a mucoid polymer and a biosurfactant. The biosynthetic genes of the mucoid polymer were previously described [42], and we identify the biosynthetic genes of the biosurfactant in this study. Competition and microscopy analyses of the extracellular polysaccharide and biosurfactant mutants reveal that these extracellular secretions function collectively to confer fitness benefit as a direct result of the spatial structures they form.

## METHODS

### Strains and culture conditions

Liquid and solid Lennox LB media (Fisher) were used for general overnight cultures. *Pseudomonas* Agar F (PAF) media (Difco) was used for all phenotypic screens, competitions, and microscopy. *Pseudomonas* minimum medium (PMM: 3.5mM Potassium phosphate dibasic trihydrate,2.2mM potassium phosphate monobasic, 0.8mM ammonium sulfate, 100mM Magnesium sulfate, 100mM sodium succinate) was used to selectively grow *P. fluorescens* isolates from conjugations with *Escherichia coli* donors. Routine cloning was carried out in *E. coli* 10B (Invitrogen) or *E. coli* JM109 (Promega), and *E. coli* S17.1λpir [43] was used as the donor strain in conjugations. When required, antibiotics were added to the media at the following final concentrations: kanamycin (50µg/mL), streptomycin (50µg/mL), ampicillin (100µg/mL), and gentamicin (20µg/mL). *P. fluorescens* was cultured at 30°C or at room temperature (~22°C), and *E. coli* strains were cultured at 37°C. Liquid cultures were incubated while shaking at 250 rpm. All *P. fluorescens* strains used in this study are listed in Table 1.

**Table 1.** *P. fluorescens* strains used in this study.

| Strain | Relevant genotype | Relevant phenotypes | Source |
| --- | --- | --- | --- |
| Pf0-1 | WT | Non-mucoid, no biosurfactant | [63] |
| Pf0-1S | WT (*Tn7-Sm<sup>R</sup>*) | Streptomycin resistance | [23] |
| Pf0-1R | WT (*Tn7-DsRed-Express*) | Red fluorescence | [23] |
| $\Delta rsmE$ | WT ($\Delta Pfl01\_1912$) | Mucoid, biosurfactant | This study |
| $\Delta rsmA$ | WT ($\Delta Pfl01\_4273$) | Non-mucoid, no biosurfactant | This study |
| $\Delta rsmI$ | WT ($\Delta Pfl01\_4104$) | Non-mucoid, no biosurfactant | This study |
| M | WT (126<sup>th</sup> nucleotide deleted in *rsmE*) | Mucoid, biosurfactant | [23] |
| MK | M (*Tn7-Km<sup>R</sup>*) | Kanamycin resistance | [23] |
| MG | M (*Tn7-Gfpmut2*) | Green fluorescence | [23] |
| M* | M ($\Delta Pfl01\_3834$) | Non-mucoid, biosurfactant | [42] |
| M*K | M* (*Tn7-Km<sup>R</sup>*) | Kanamycin resistance | This study |
| M*G | M* (*Tn7-Gfpmut2*) | Green fluorescence | [42] |
| M<sup>S</sup> | M ($\Delta Pfl01\_2211$) | Mucoid, no biosurfactant | This study |
| M<sup>S</sup>K | M<sup>S</sup> (*Tn7-Km<sup>R</sup>*) | Kanamycin resistance | This study |
| M<sup>S</sup>G | M<sup>S</sup> (*Tn7-Gfpmut2*) | Green fluorescence | This study |
| M<sup>S</sup>* | M ($\Delta Pfl01\_2211 \Delta Pfl01\_3834$) | Non-mucoid, no biosurfactant | This study |
| M<sup>S</sup>*K | M<sup>S</sup>* (*Tn7-Km<sup>R</sup>*) | Kanamycin resistance | This study |
| M<sup>S</sup>*G | M<sup>S</sup>* (*Tn7-Gfpmut2*) | Green fluorescence | This study |

### Quantitative PCR

Total RNA was isolated from colonies grown for 3 days at room temperature on PAF plates using the TRIzol® Reagent (ThermoFisher) under the manufacturer’s protocol. Total RNA quality and concentration were assessed using the NanoDrop spectrometer. First strand cDNA synthesis was carried out using the High-Capacity RNA-to-cDNA kit (Applied Biosystems) with 1µg of RNA and random hexamers following the manufacturers protocol. qPCR optimized primers were obtained from Integrated DNA Technologies (Table 2) and their quality was assessed through PCR with gDNA, cDNA, and no-RT cDNA reactions. qPCR was performed using SYBR Green (ThermoFisher) on the StepOnePlus™ instrument (Applied Biosystems). Each reaction was analyzed to ensure only one amplicon was amplified using dissociation curves. Gene expression was calculated using the 2^−ΔΔCT^ method with the 16S rRNA gene as an internal reference and quantified relative to *rsmI* expression [44].

**Table 2.**
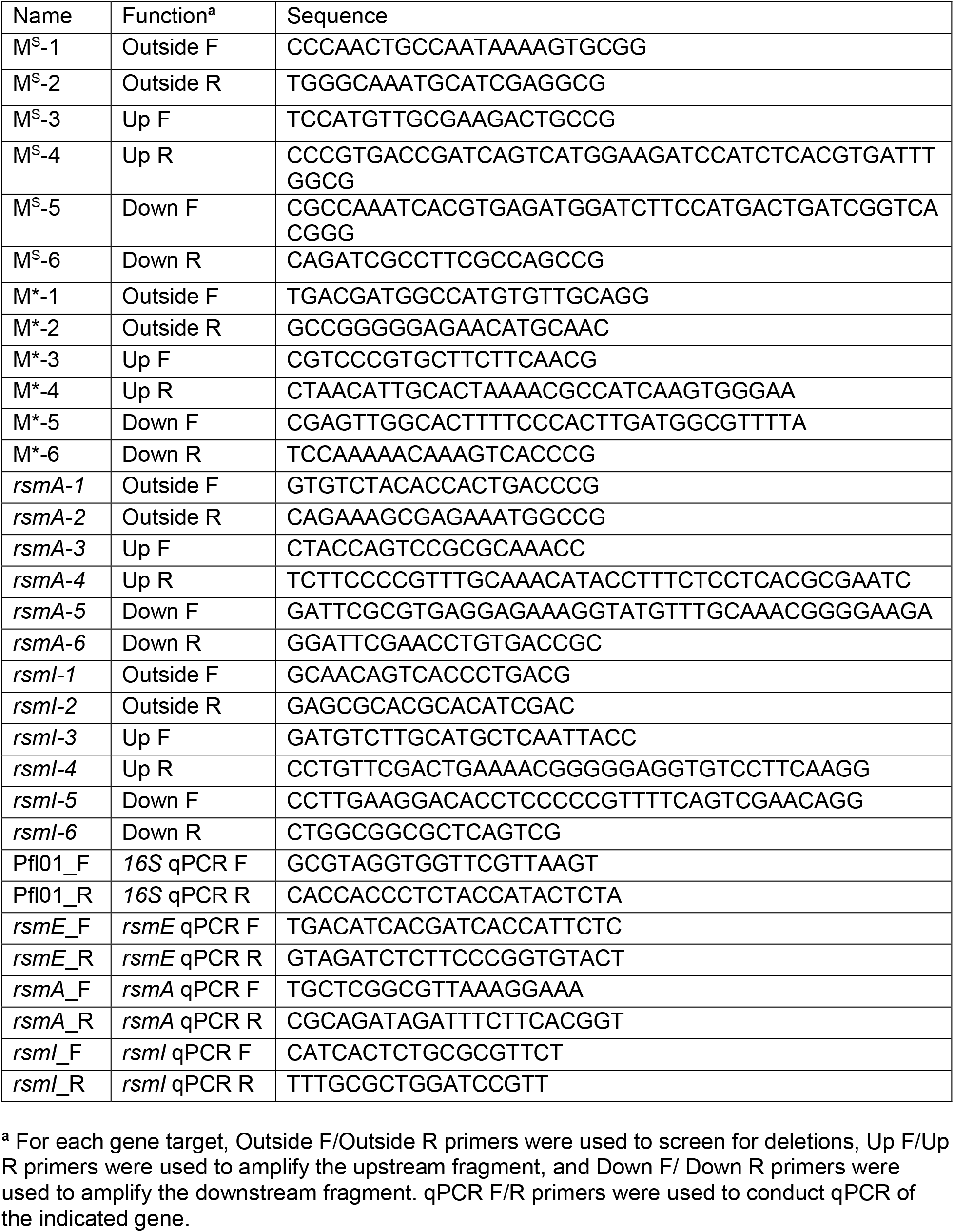
Primers used in this study.

### Biosurfactant assay

Nuclepore Track-Etch polycarbonate membranes (Whatman: 0.4µM pore size, 90mm diameter) were used for assessing biosurfactant production. As previously described [23], one side of the membrane is shiny and the other is dull due to the manufacturing process. The dull side’s surface contains gaps and ridges that physically trap cells, but the biosurfactant permeates to produce the visible ring around colonies. The shiny side’s surface is smooth, which allows biosurfactant producing cells to spread out through growth. Sterile forceps were used to overlay the membrane on the PAF agar surface, and 20µl of overnight culture was spotted directly on the membrane and allowed to fully dry before the plates were inverted and incubated over night at room temperature.

### Identification of the biosurfactant biosynthesis genes by transposon mutagenesis

Random transposon mutagenesis, using the plasmid pUT-mini*T*n5-Km*lacZ2* [45, 46] in *E. coli* S17.1λpir as the donor, was carried to identify the biosynthesis genes of the biosurfactant as previously described to identify the biosynthesis genes of the mucoid polymer [42]. Briefly, overnight cultures of the donor and M* strains were washed in PMM, mixed at the relative ratio of 1:6, spotted on solid LB to conjugate, incubated at 30°C for three hours, harvested and plated out on solid PMM supplemented with kanamycin. Over 20,000 transconjugant colonies were picked and rearrayed using the QBot (Genetix) into 384-well plates containing kanamycin supplemented PMM, then incubated at 30°C. Surfactant assays on overnight cultures were conducted on PMM plates overlayed with the dull side of the polycarbonate membrane as described above with a disposable 384 pin replicator (Scinomix). Mutants that were defective in biosurfactant production (dull side) were rearrayed into 96-well plates containing kanamycin supplemented PMM, then incubated at 30°C. Overnight cultures were retested for biosurfactant production as described above using a disposable 96 pin replicator (Scinomix). Mutants that failed to produce the biosurfactant ring were selected, ignoring ones that had obvious growth defects. The transposon insertion sites were identified by arbitrary primed PCR as previously described [42].

### Mutant construction and tagging

Gene deletion mutants were constructed by the gene splicing by overlap extension method [47], using the plasmid pMQ30[48] or pSR47s [49] as previously outlined [23, 42]. PCR primers used to construct and confirm each mutation are listed in Table 2. Briefly, for each targeted gene, approximately 500bp of its flanking upstream and downstream regions were individually amplified, joined together, first cloned into the pGEM-T Easy Vector system (Promega) then sub-cloned into pMQ30 or pSR47s, and transformed into *E. coli* S17.1λpir as the donor strain. Overnight cultures of the donor and target strains were washed in PMM and mixed at an equal ratio, spotted on solid LB, incubated at 30°C overnight, harvested and plated out on solid PMM supplemented with gentamicin (pMQ30) or kanamycin (pSR47s). Transformants were grown on solid LB supplemented with sucrose (5%, w/v) overnight, and the resulting colonies were screened using primers that bind outside the two flanking fragments for expected reduction in the amplicon size. To confirm the gene deletions, we isolated genomic DNA from overnight cultures using the DNeasy UltraClean Microbial Kit (Qiagen) following the manufacturer’s protocol and whole genome sequencing was conducted at the Microbial Genome Sequencing Center (MiGS; Pittsburgh, PA). Kanamycin-resistant and streptomycin-resistant strains used in competitions and GFP-tagged and DsRed-Express-tagged strains used in microscopy were constructed using the mini-*Tn7* chromosomal insertion system [50] as previously described [23, 42].

### Measurement of monoculture growth

For the measurement of growth in colonies, overnight cultures were resuspended in PMM and 20µL was spotted on PAF plates and incubated at room temperature. To enumerate the initial population size, each cell suspension in PMM was serially diluted and plated out on LB plates, and resulting colonies were counted on the following day. Three spotted colonies were scraped on each day over seven days and resuspended in 5 mL of PMM using a sterilized bent glass Pasteur pipette. Cell suspensions were vortexed until clumps were no longer visible then serially diluted and enumerated as above. For the measurement of growth in liquid, overnight cultures were diluted into a manually formulated PAF without agar [51] in six replicates, and optical density at 600 nm was measured every 30 minutes over 48 hours (30°C, constant shaking) in the Bioscreen C MBR (Oy Growth Curves Ab Ltd.).

### Competition assay

Competitions between kanamycin-resistant mutant strains and streptomycin-resistant WT strain were conducted as previously described [23]. Briefly, overnight cultures (1.5 mL) were washed in fresh PMM and re-suspended in 1.0 mL PMM, and the mutant strain suspension was serially diluted to 10^−3^ in PMM and mixed with equal volumes of the undiluted WT strain suspension. 20µL of each competition mixture was spotted in triplicate on a PAF plate and incubated at room temperature. To enumerate the initial population size of the competing strains, each competition mixture was serially diluted and plated out on LB plates supplemented with either kanamycin or streptomycin, and resulting colonies were counted on the following day. Four or seven days later, the spotted colonies were scraped and resuspended in 5mL of PMM, serially diluted, plated, and counted as for the initial competition mixture. The results of the competitions were analyzed by calculating the relative fitness (*W*) of each competing strain against the WT [52].

### Statistical analysis

Competition experiments were conducted with at least three biological replicates and two technical replicates for each biological replicate. The data were first analyzed with ANOVA to evaluate if the means of the biological replicates differ significantly, then Tukey’s honest significant difference test (p < 0.05) was applied to make multiple pairwise comparisons within the dataset. All comparisons were found to be statistically different or noted as n.s. otherwise. Statistical tests were conducted using GraphPad Prism.

### Microscopy

Overnight cultures of GFP-labeled strains and DsRed-Express-labeled WT were washed and resuspended in PMM. All GFP-labeled cell suspensions were serially diluted to 10^−5^ in PMM and mixed with equal volumes of the undiluted DsRed-Express-labeled WT suspension. 20µL of each competition mixture was spotted in triplicate on PAF plates and incubated at room temperature. Epifluorescence microscopy was conducted using the Nikon SMZ25 stereo-compound microscope with the 0.5 X SHR Plan Apo objective and the NIS Elements software. For confocal microscopy, an agar slice containing the entire colony was placed on a microscope slide and visualized without a coverslip. Confocal microscopy was conducted using the Nikon Ti2 microscope with the 20X TU Plan Fluor objective or the air-corrected 100X TU Plan Apo objective and the NIS Elements software. Non-fluorescent imaging of colonies was carried out using the Hayear overhead microscope (HY-2307) or the Canon Rebel EOS T3 DSLR camera. Images were rendered using the NIS Elements and ImageJ software.

## RESULTS

### RsmE, RsmA, and RsmI in *Pseudomonas fluorescens* Pf0-1 are highly conserved in sequence and all three respective genes are simultaneously expressed

*Pseudomonas fluorescens* Pf0-1 possesses three Rsm paralogs – RsmA, RsmE, and RsmI – that share high sequence similarity (Fig. 1A). We sought to first determine whether or not all three respective genes are expressed. Quantitative PCR confirmed that all three genes are indeed simultaneously expressed, with *rsmA* and *rsmI* transcripts being the most and least abundant, respectively (Fig. 1B). These results show that the exclusive selection of *rsmE* mutations in our previous experimental evolution study [23] was not simply due to the absence of *rsmA* and *rsmI* expression under the same experimental conditions.

**Fig. 1.**
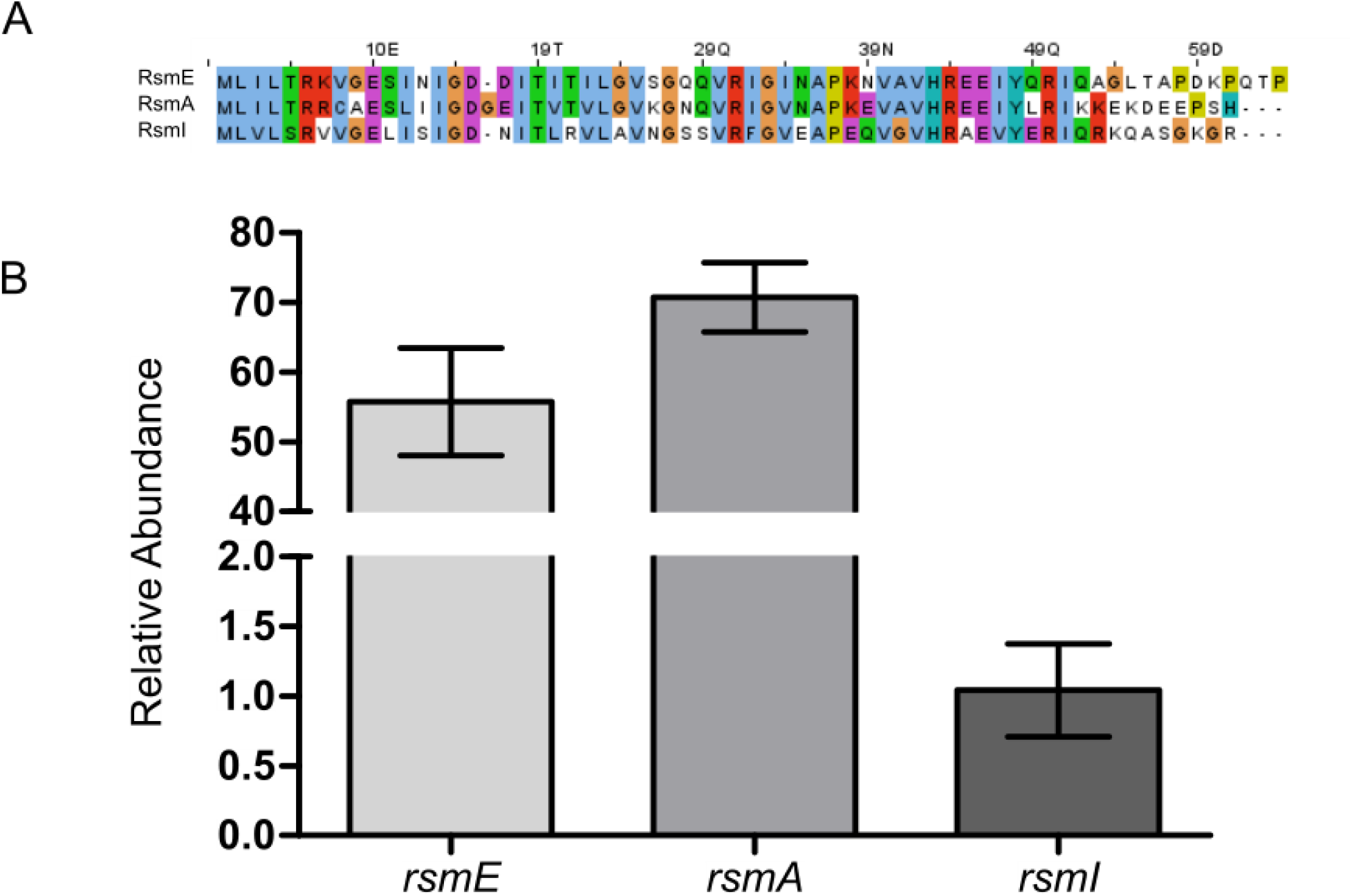
Rsm paralogs in *P. fluorescens* Pf0-1 share a highly conserved sequence and their respective genes are simultaneously expressed. (A) Sequence alignment of Rsm-paralogs in *Pseudomonas fluorescens* Pf0-1 with ClustalX show similarities of amino acid sequence and chemical properties [62]. (B) Expression of *rsmE, rsmA*, and *rsmI* genes assessed in WT by qPCR. Transcripts of all three genes were detected and shown here is the relative abundance of each transcript using the 2^-ΔΔCT^ method compared to that of the least abundantly expressed *rsmI*. Plotted are the mean of three biological replicates with three technical replicates for each biological replicate, and the error bars represent the standard deviation of the mean.

### RsmE specifically regulates the production of a mucoid polymer and biosurfactant

Experimentally selected *rsmE* mutants visibly produce a mucoid polymer and/or a biosurfactant [23], which suggests that specific mutations differentially impact RsmE’s function. To determine if these extracellular secretions are commonly regulated by the three Rsm homologs, we constructed deletion mutants of the respective genes. Comparison of colony morphologies show that only the *rsmE* mutant exhibits mucoidy (Fig. 2A). In addition, mucoid patches consistently emerge in colonies of WT, *rsmA* mutant, and *rsmI* mutant (Fig. 2A), which are characteristic of naturally mutated *rsmE* [23]. These results confirm that the production of the mucoid polymer is specifically regulated by RsmE. We next compared biosurfactant production on a polycarbonate membrane overlaid on the agar surface. Production of the biosurfactant on the shiny side of the membrane allows the colony to spread out radially, but the cells remain trapped on the dull side of the membrane while the biosurfactant spreads out unhindered [23]. Only the *rsmE* mutant produced a visible ring on the dull side of the membrane and also spread out on the shiny side of the membrane (Fig. 2B). These results confirm that the production of both the mucoid polymer and the biosurfactant is uniquely governed by RsmE from its paralogs.

**Fig. 2.**
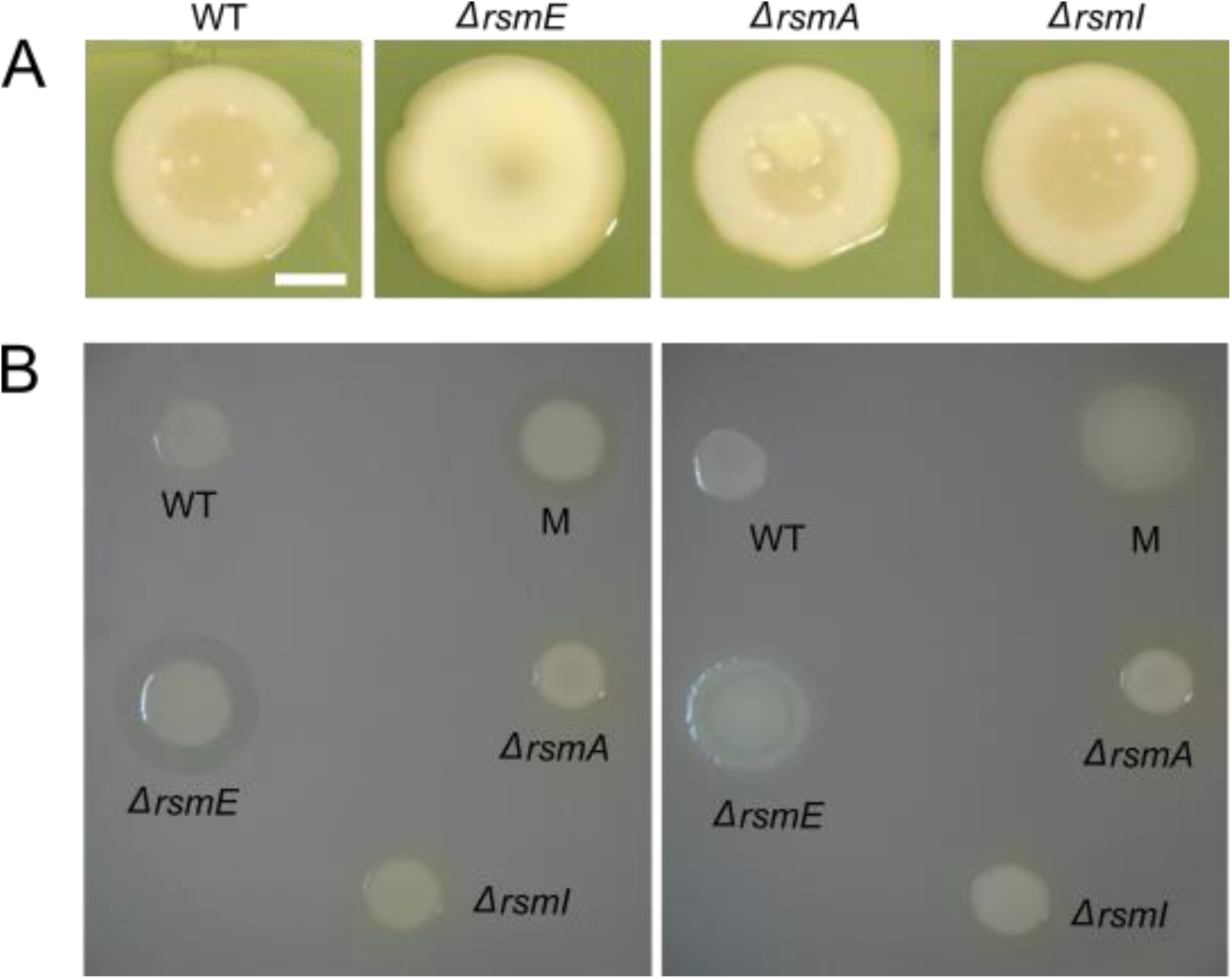
Both the mucoid polymer and the biosurfactant are regulated by RsmE, but not by RsmA or RsmI. (A) Colony morphology comparisons of WT and deletion mutants of *rsmE, rsmA*, and *rsmI*. Liquid cultures were spotted on PAF plates seven days prior to capturing the images. Only the *ΔrsmE* strain is mucoid in appearance and new mucoid patches naturally emerge in WT, Δ*rsmA*, and Δ*rsmI* colonies that characteristically represent de novo *rsmE* mutations. The scale bar represents 10 mm. (B) Comparison of biosurfactant production on the dull (left) and shiny (right) sides of the polycarbonate membrane overlaid on PAF. The M strain is a naturally selected mutant from a WT colony, harboring a frameshift mutation in *rsmE*. Only the Δ*rsmE* and M strains produce the biosurfactant ring on the dull side that promotes spreading of cells on the shiny side of the membrane.

### Identification of the biosurfactant as gacamide A

The biosynthetic genes of the mucoid polymer were previously characterized to encode a glucose-rich extracellular polysaccharide, and a corresponding gene was deleted in a mucoid (M) strain with a frameshift mutation in *rsmE* [23] to produce the non-mucoid M* strain [42]. To identify the biosynthetic genes of the biosurfactant, we carried out a random transposon mutagenesis in the M* strain background. Seven mutants were independently isolated that no longer produced the secretion on the dull side of the polycarbonate membrane and failed to spread out on the shiny side of the membrane. All transposon insertion sites were mapped to three contiguous loci (annotated as Pfl01_2211, Pfl01_2212, and Pfl-1_2213), which were recently demonstrated to encode non-ribosomal peptide synthetases [53] that produce the cyclic lipopeptide gacamide A [54]. Cyclic lipopeptides are indeed classified as a surfactant, and they contribute to surface motility and biofilm formation in many *Pseudomonas* spp. [55, 56]. Given that four independent transposon insertions occurred in the Pfl01_2211 locus, we constructed a corresponding in-frame deletion mutant in the M strain to produce the M^s^ strain, and the same mutation was also introduced in the non-mucoid M* strain to produce the M^S^* strain. Neither M^S^ nor M^S^* produce the biosurfactant ring on the dull side of the membrane and the spreading phenotype on the shiny side of the membrane (Fig. 3), confirming that the Pfl01_2211-Pfl-1_2213 cluster encodes the production of the biosurfactant. Importantly, M* maintains the production of the biosurfactant and M^S^ maintains the production of the mucoid polymer (Fig. 3), which shows that the biosynthesis of these two secreted products are not genetically linked to one another, but are both regulated by RsmE.

**Fig. 3.**
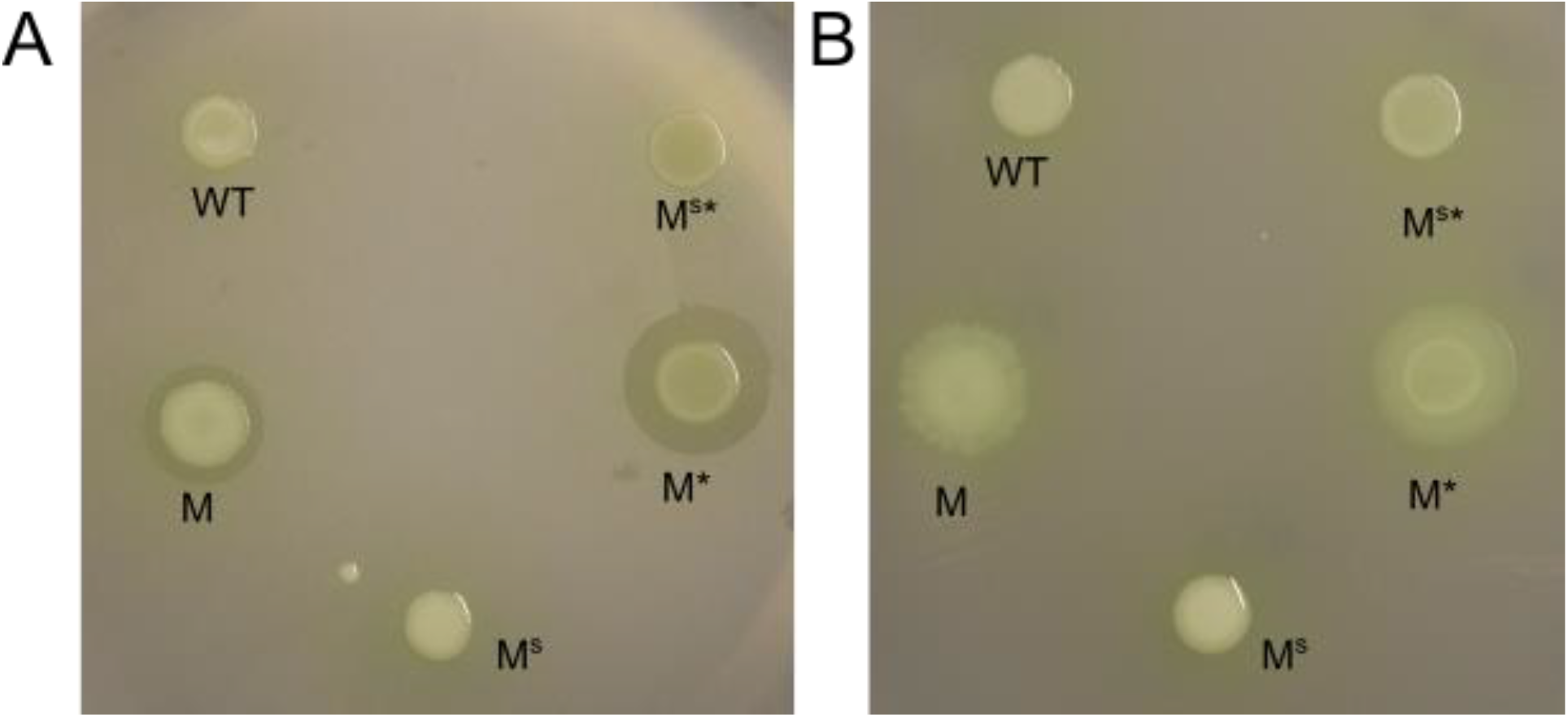
Deletion of the Pfl01_2211 locus abolishes biosurfactant production. Shown are the results from the dull side (A) and the shiny side (B) of the polycarbonate membrane. M (*rsmE* mutant) and M* (M with the mucoid polymer biosynthesis gene (Pfl01_3834) deleted) produce the biosurfactant and spread on the surface, but M^S^ (M with Pfl01_2211 deleted) and M*^S^ (M with both Pfl01_3834 and Pfl01_2211 deleted) fail to do so like the WT with an unaltered *rsmE* gene. These results confirm that the Pfl01_2211-2213 cluster encodes the biosynthetic genes of the biosurfactant, which is now known to be gacamide A.

### Both the mucoid polymer and biosurfactant confer competitive advantage

All experimentally selected *rsmE* mutants outcompete the WT strain in co-cultured colonies [23]. To assess the contributions of the RsmE-regulated mucoid polymer and the biosurfactant, we independently competed M, M^S^, M*, and M^S^* against the WT in co-cultured colonies and assessed their fitness relative to the WT. All four strains outcompeted the WT throughout the duration of the experiments (Fig. 4), with M and M^S^ being nearly equal in fitness and M* and M^S^* exhibiting decreased fitness at day 4. However, we observed reduced fitness in all secretion mutants compared to M by day 7, with M^S^ and M* being comparable and M^S^* exhibiting further reduction. Such step-wise decreases in fitness indicate that each secreted product independently confers competitive advantage and the two secretions also likely function in an additive manner. Furthermore, the fact that M^S^* retains the ability to outcompete the WT indicates that there are additional RsmE regulated genes that contribute to M’s dominance over the WT.

**Fig. 4.**
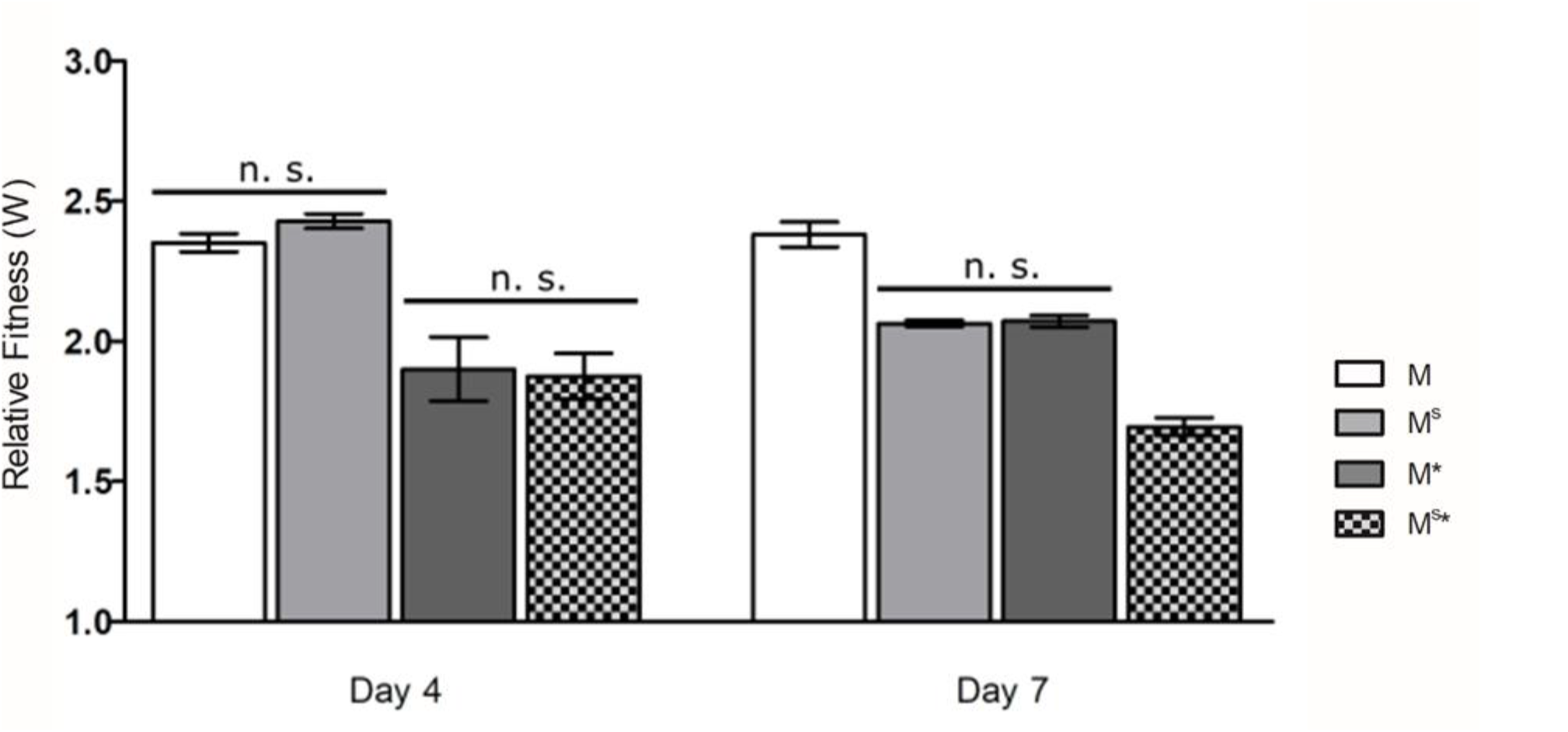
Competitions of M, with or without mucoid polymer and/or biosurfactant production, against WT show varying levels of relative fitness over time. WT was chromosomally tagged with streptomycin resistance and all mutants were tagged with kanamycin resistance, and these resistance markers produce neutral relative fitness in *P. fluorescens* Pf0-1 [23]. Error bars represent the standard deviation of the mean relative fitness (mutant over WT) calculated from three independent populations after four and seven days of incubation. Dataset from each time point was analyzed by ANOVA (p < 0.0001) and Tukey’s honest significant difference test showed that all pairwise comparisons were significantly different (p < 0.05) except for those indicated as nonsignificant (n. s.). Relative fitness (W) of 1 indicates equal fitness of the mutant and WT and a W of greater than 1 indicates that the mutant outcompeted the WT. Both the mucoid polymer and the biosurfactant provide competitive advantage. However, M^S^* (*rsmE* mutant with biosynthesis genes of both secretions deleted) still outcompetes the WT, suggesting that additional RsmE-regulated products contribute to the competitive advantage of M (*rsmE* mutant) against the WT.

### The mucoid polymer creates space and the biosurfactant prevents the diffusion of the mucoid polymer at the colony surface

The temporal differences in the relative fitness between M^S^ and M* (Fig. 4) suggests that the mucoid polymer plays a more significant role early in the competition. Importantly, our secretion mutants exhibit equal growth profiles compared to the WT as monoculture in both liquid and colonies (Fig. 5). The M data here recapitulates the results from our previous study, which also demonstrated that the competitive advantage of *rsmE* mutants specifically requires the formation of spatial structures that decreases local density and provides greater access to oxygen [23].

**Fig. 5.**
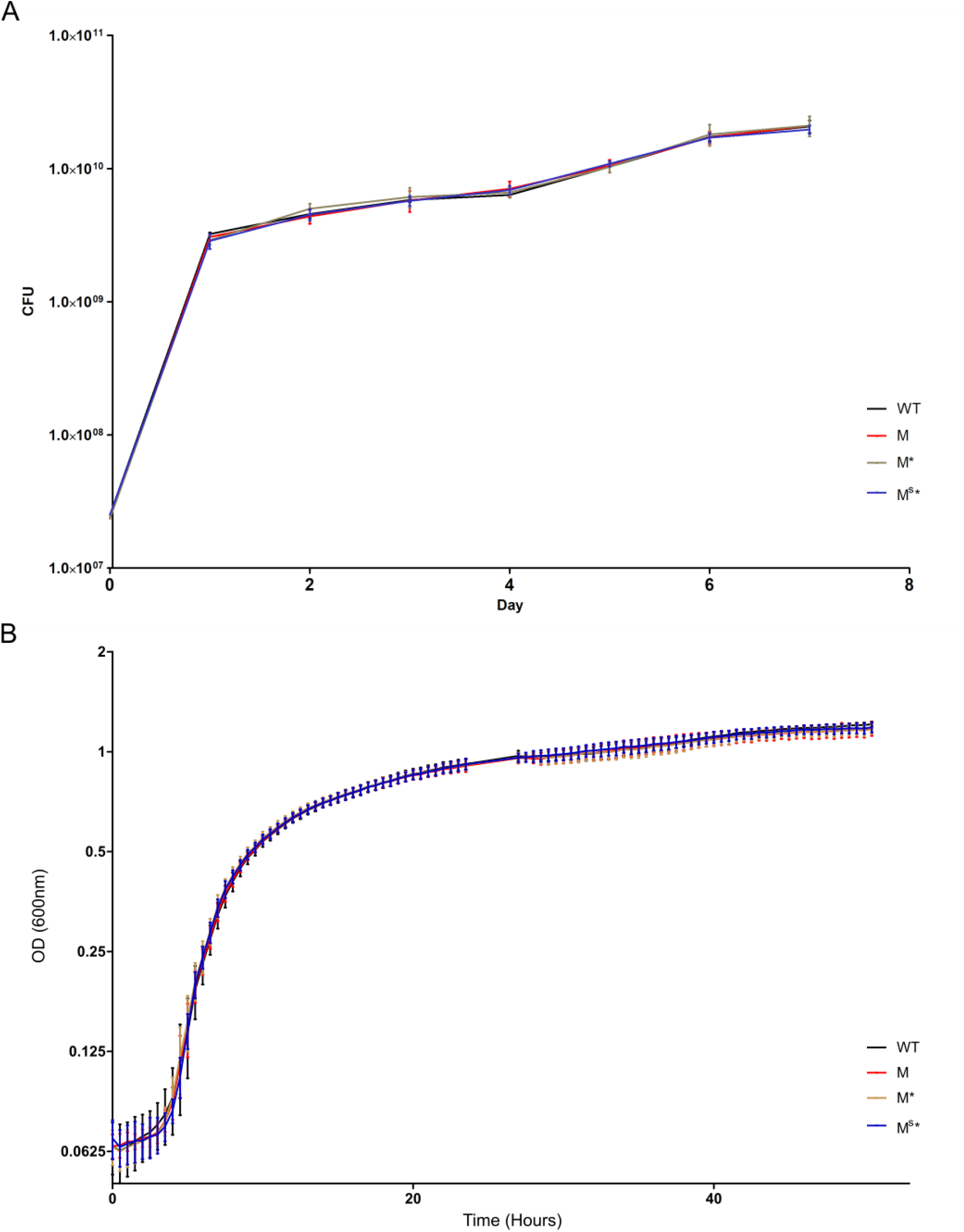
Production of RsmE-regulated extracellular secretions does not impact growth in monoculture. (A) Growth profiles of single genotype colonies of WT, M, M*, and M^S^* on solid PAF. Each data point represents the mean colony forming units (CFU) of three populations and the error bars represent the standard deviation of the mean. (B) Growth profiles of single genotypes in liquid PAF as measured by optical density at 600 nm. Shown are the mean of six independent cultures for each strain, and the error bars represent the 95% confidence interval.

To explore the functional role of the RsmE-regulated mucoid polymer and biosurfactant in spatial structure formation, we carried out epifluorescence and confocal microscopy analyses of our collection of secretion mutants against the WT. We first introduced a constitutively expressed *gfp* gene into the chromosome of WT, M, M^S^, M*, and M^S^* strains. Each green fluorescent strain was mixed with red fluorescence labeled WT at the respective ratio of 10^−5^:1 to best visualize isolated spatiogenetic structures in colonies after 5 days. Epifluorescence imaging of entire colonies shows isolated green fluorescent patches emerging from mostly red fluorescent WT colonies, with M and M^S^ producing consistently bigger patches compared to M* and M^S^* (Fig. 6A). Each co-culture also produced red fluorescent mucoid patches, which represents *de novo rsmE* mutants naturally emerging from the red fluorescent WT cells [23], however, no green fluorescent patches were observed in the WT:WT colonies. With confocal imaging at a low magnification, the green fluorescence signal in the smaller patches formed by M* and M^S^* is much more intense compared to those formed by M, and M^S^ patches produced the least intense fluorescence signal (Fig. 6B). Individual patches formed by both M and M^S^ typically merged together with nearby patches through continuous expansion over time, but we consistently observed M^S^ patches to be much more amorphous in structure with less defined individual boundaries. In contrast, green fluorescent WT patches were rarely observed and appeared to comprise only few cells.

**Figure. 6.**
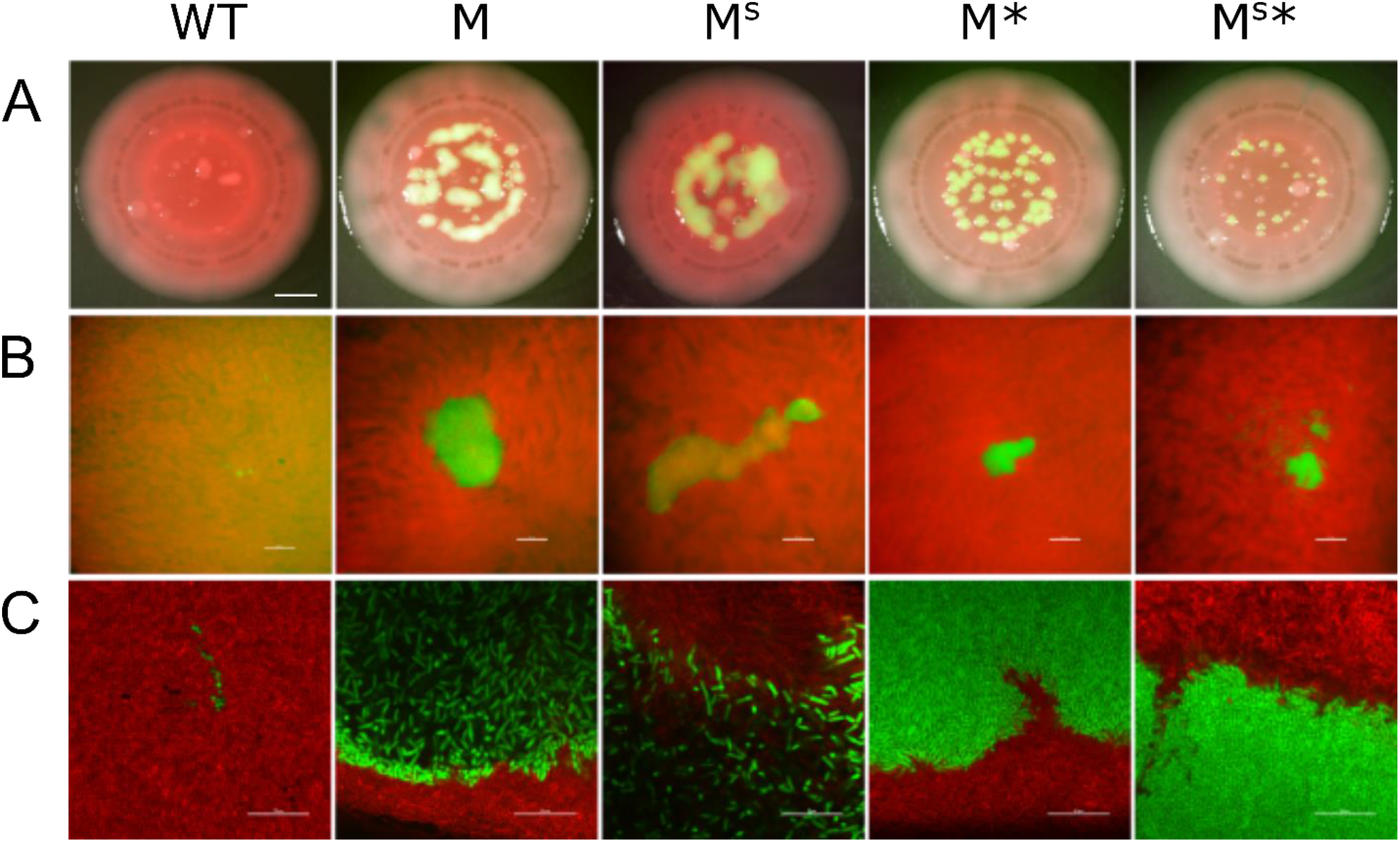
The mucoid polymer and biosurfactant function together in the formation of a dominant spatial structure. Each indicated strain was chromosomally tagged with GFP, heavily under-represented in a mixture with DsRed-Express-tagged WT, and representative co-cultured colonies were imaged five days later. (A) Epifluorescence microscopy images that capture the entire colony (A, scale bar represents 10 mm). Each sample shows the natural emergence of red mucoid patches that are characteristic of *de novo rsmE* mutants stemming from the red-fluorescent WT cells. (B) Confocal microscopy images focusing on the surface of individual patches at a low magnification (scale bar represents 50 um). M^S^* produces unique patches that appear to be mixed with red WT cells. (C) Confocal microscopy images at a higher magnification focusing on the boundaries between the mutant and WT (scale bar represents 10 um). The mucoid polymer is solely responsible for creating the space of low cell density (black space is devoid of cells) and the biosurfactant appears to physically hold the mucoid polymer and producing cells from flowing out from the newly created space. M^S^* produces the smallest patches that are densely filled, as reflected by vertically aligned cells (spheres) similar to the WT:WT spatial organization (left panel). However, M^S^* maintains the ability to form an organized structure that excludes WT cells, suggesting that additional RsmE-regulated products contribute to the spatial dominance of M.

Confocal imaging using an air-corrected 100X Plan Apo objective provided a clear view of individual green fluorescent cells and their spatial arrangement within a given patch surrounded by red fluorescent WT cells (Fig. 6C). M cells were present at a strikingly lower density compared to the neighboring WT cells, with the characteristic black space that is devoid of cells [23]. In addition, M patches are defined by a clear boundary formed with a thin layer of M cells which appears to exclude the encroachment of WT cells into the black space. In contrast, M^S^ patches lacked a clear exclusionary boundary, with M^S^ cells appearing to flow over the WT cells. This interpretation is also reflected in the lower magnification observations of M^S^ patches being more mucoid and amorphous (Fig. 6A) and producing less intense fluorescent signal (Fig. 6B) compared to patches formed by M. M* and M^S^* both formed much densely packed patches with clear boundaries against the WT cells, but M^S^* cells appear to be even more packed as indicated by the uniquely vertical arrangement of cells (Fig. 6C) and much smaller size of individual patches (Fig. 6A). These observations collectively suggest that the mucoid polymer is the primary driver of creating space while the biosurfactant spatially sequesters the mucoid polymer to prevent their diffusion. However, M^S^* retains the ability to produce a spatiogenetic structure that contrasts greatly from the green fluorescent WT cells that form small clusters of only few cells without any organized structure (Fig. 6C), likely representing daughter cells stemming from initially a single mother cell. As reflected by our relative fitness data (Fig. 4), there appears to be additional RsmE-regulated genes that specifically promote spatial competition in a densely populated colony.

## DISCUSSION

Several members of the gamma-proteobacteria, including *Pseudomonas* spp., possess multiple paralogs of CsrA/Rsm proteins, and their corresponding genes are also present in diverse plasmids and bacteriophages [27]. We had previously shown that mutations in *rsmE* are repeatedly selected as mucoid patches in colonies of *P. fluorescens* Pf0-1 by creating space and capturing optimal positioning within a crowded environment [23]. The exclusive association of *rsmE* mutations with this striking phenotype suggests that RsmE’s function is not entirely redundant from that of its paralogs, RsmA and RsmI. In this study, we show that all three paralogs are accessible to evolutionary selection, since their respective genes are simultaneously expressed under the same experimental conditions. Furthermore, mucoid patches consistently emerged in both *rsmA* and *rsmI* knockout colonies, much like those that emerge from WT colonies through *de novo* mutations in *rsmE*. These observations strongly support our prediction that the formation of beneficial spatial structures occurs specifically through mutations that deregulate RsmE’s native function.

We have shown that knocking out *rsmE* results in the production of two visible extracellular secretions, a mucoid polymer and a biosurfactant, but neither are produced in *rsmA* nor *rsmI* knockouts. Thus, RsmE appears to either directly repress the production of these secretions or modulate the activity of other regulators that directly govern their production. Genetically removing the production of either or both secretions in the *rsmE* mutant significantly reduced competitive advantage against WT in co-cultured colonies. However, all engineered secretion mutants shared the same growth profiles compared to WT in liquid and colony monocultures. These observations collectively suggest that both secretions contribute to the spatial structure formation by the *rsmE* mutant, and we confirmed this prediction through epifluorescence and confocal microscopy.

The two key characteristics associated with the dominant spatial structure formed by the *rsmE* mutant are creation of space with low cellular density and exclusion of the neighboring WT cells from this local environment [23]. Here, we have demonstrated that the mucoid polymer is solely responsible for creating the space. We had initially interpreted that the biosurfactant forms the exclusionary boundary due to the mixed presence of the biosurfactant knockout and WT cells. However, we consistently observed that the WT cells rarely invade deeply into the areas of low cellular density at high optical magnification. In addition, the borders of individual patches formed by the biosurfactant mutant were less defined and the mutant cells appeared to flow out on top of the neighboring WT cells, akin to outflowing lava from a volcano. However, these observations indirectly contradict the results of our membrane assay, which showed that the same biosurfactant promotes the spreading of cells on the membrane surface. In fact, we initially referred to the corresponding secretion as a biosurfactant, due to the well-known function of bacterial surfactants that reduce surface tension to promote swarming on semi-solid agar surfaces [57].

We identified the biosynthetic genes of the biosurfactant in this study, which were recently characterized by an independent group to produce a cyclic lipopeptide named gacamide A that promotes swarming [54]. *Pseudomonas* spp. produce numerous cyclic lipopeptides that variably contribute to surface-spreading and biofilm formation, and this variability potentially depends on discrete interactions with diverse extracellular or cell membrane-associated products [56, 58]. The amphiphilic structure of gacamide A likely promotes its interaction with both hydrophilic compounds, like the mucoid polymer, and hydrophobic compounds that co-accumulate within the patches formed by the *rsmE* mutant. Importantly, removing the production of both the mucoid polymer and gacamide A maintained the respective *rsmE* mutant’s ability to outcompete the WT, albeit with much reduced spatial dominance. These observations suggest that there are additional RsmE-regulated products that contribute to the competitive advantage of the *rsmE* mutant, which clearly manifests through beneficial structures [23]. Pressure likely builds up internally within a localized patch as the accumulating mucoid polymer constantly pushes away the surrounding WT cells to expand space. We thus speculate that gacamide A physically stabilizes the mucoid polymer and additional RsmE-regulated products to prevent their diffusion at the surface of the colony, which is uniquely devoid of neighboring cells and provides much less resistance.

A potential criticism of this study is the utilization of bacterial colonies to explore spatial structure formation, which lack important mechanical properties that manifest in natural microbial communities [7]. However, resolving the problem of space and resource constraints in a densely populated colony likely shares common principles with other organisms in different experimental systems. Extracellular polysaccharides produced by *Vibrio cholerae* growing in microfluidic device biofilms promote the formation of isogenic structures that exclude the neighboring non-producers [59], and glycolipid biosurfactants produced by *Streptococcus* spp. selectively displace competing genotypes on the tooth surface [60]. Cyclic lipopeptide production in *Bacillus subtilis* is essential for fruiting body formation on an agar surface, and mutants that lack this biosurfactant initially form projecting columns, but they grow laterally and subsequently fuse together [61], much like our biosurfactant knockout cells. Our study also establishes a highly tractable experimental pipeline to identify and characterize additional RsmE-regulated products, and to explore why RsmA and RsmI are functionally excluded from the formation of spatial structures.

## ACKNOWLEDGEMENTS

This study was funded by the National Institute of General Medical Sciences of the NIH 1R15GM132856 (W.K.).

## AUTHOR CONTRIBUTIONS

W.K. designed the study, A.E., J.D., W.M., R.S., A.D., and W.K. performed experiments, A.E., M.W., J.D., and W.K. analyzed data, and A.E., M.W., and W.K. wrote the manuscript.

